# Functional insights on chemical communication of fishing cats: a strategic mechanism for adaptation in semi-aquatic habitat

**DOI:** 10.1101/2024.11.19.624236

**Authors:** Subhadeep Das, Payel Das, Sourav Manna, Mousumi Poddar Sarkar

**Author notes:** email id.

## Abstract

Chemical communication system of fishing cats, focusing on volatile organic compounds present in their urine (UR) and ‘marking fluid’ (MF) has been explored. Using solid-phase microextraction guided gas chromatography mass spectrometry analysis, a diverse range of functional groups from UR and MF were detected, including ketones, aldehydes, sulfur- and nitrogen-containing compounds, among which 3-Mercapto-3-methylbutanol (3MMB) and its related sulfur compounds were found to be sex-specific, predominantly present in males. The unique behavior of fishing cats to urinate frequently in water when experimentally tested with synthetic 3MMB, was aligned for its retention in aqueous medium upto 21 days. This suggests that fishing cats may employ an energy-efficient scent-marking strategy by leveraging aquatic environments for prolonging chemical signalling process. The analysis of less-volatile compounds, such as triglycerides and fatty acids, suggests that these compounds could be acted as fixatives, helping to stabilize and prolong the intensity of chemical messages coded to highly volatile molecules. The combination of highly volatile, less volatile compounds and lipids may create complex synergy to facilitate interindividual communication, potentially serving as pheromones. This research enhances our understanding of the fishing cat’s adaptation to semi-aquatic environments and its unique behavioural pattern in marshy landforms, thus concentrated with high population density in deltaic South Bengal.

## Introduction

The fishing cat [*Prionailurus viverrinus* (Bennett, 1833)], a medium-sized wild cat species, native to South and South-East Asia, is uniquely adapted to wetland ecosystems of South Bengal (Mukherjee et al. 2016). With its semi-aquatic lifestyle, this felid acquires a wide range of physiological and behavioural attributes that facilitate its survival in riparian habitats, including specialized hunting techniques and with a distinctive diet, primarily comprised of fishes and other aquatic prey. Despite ecological importance for its cryptic nature, the fishing cat (FC) remains one of the least understood members of the Prionailurus genus, particularly related to its communication strategies. From the historical note of Brongersma (1935), it was known that in 1819, Pierre-Médard Diard and Alfred Duvaucel collected specimen of fishing cat from an expedition to Singapore which is thought to be the world’s oldest museum specimen of fishing cats that was preserved by the Netherlands’ Naturalis Biodiversity Center, Leiden (RMNH) (Jusoh et al. 2022). Recently, a comprehensive report of fishing cat distribution in Nepal and their habitat suitability has been published, considering impact of different climatic factors and environmental parameters (Mishra et al. 2022). Phosri et al (2021) surveyed extensively on population of FCs in coastal wetlands and estuaries of Thailand and highlighted the possible reasons, like rapid habitat loss by progressive civilization, for extinction of this water-loving animal from main-land.

Chemical communication plays a crucial role in the social and reproductive behaviours of many felid species, serving functions such as territory marking, individual recognition, and mate attraction (Poddar-Sarkar and Brahmachary 1997, 2004, 2014; Burger et al. 2006, 2008; Brahmachary and Poddar-Sarkar 2015; Soso and Koziel 2016, 2017; Das et al. 2019, 2023a; Tommasi et al. 2021). In solitary and elusive species like FC, chemical signalling is especially significant, diminishing direct interactions between individuals and for receiving information about the sexual status of their mates. These chemical signals are conveyed through various excretions such as urine, faeces, glandular secretions from epidermal structures etc; each carrying complex information that can influence the behavior and physiology of the individuals. Like other felids, fishing cat marks its territory by urination, marking fluid (MF) spraying, cheek and body rubbing, and scat deposition to leave scent marks in surrounding environment (Mellen 1993; Das et al. 2023b).

In felids, the vomeronasal organ (VNO) and the main olfactory bulb are the key organs in neural circuitry that are involved in the detection and processing of chemical cues (Salazar et al. 1996; Poddar-Sarkar and Brahmachary 2014). These chemical cues can be species-specific and are often composed of pheromones and other semiochemicals, which provide information about an individual’s specific identity. While several researches have been conducted on chemical communication in other felids such as lion (*Panthera leo*), tiger (*Panthera tigris*), leopard (*Panthera pardus*), cheetah (*Acinonyx jubatus*), snow leopard (*Panthera uncia*), bob cat (*Lynx rufus*), and domestic cat (*Felis catus*), relatively little is known about how fishing cats adopt mechanism to dissipate chemical signals to utilize, in their aquatic and semi-aquatic environments (Mattina et al. 1991; Poddar-Sarkar and Brahmachary 1997, 2004, 2014; Burger et al. 2006; Miyazaki et al. 2006b, 2008; Poddar-Sarkar et al. 2008; Apps et al. 2014; Brahmachary and Poddar-Sarkar 2015; Soso and Koziel 2016, 2017; Das et al. 2019; Tommasi et al. 2021).

The study of chemical communication in fishing cats is particularly relevant for exploring the reasons for their vulnerability as stated by International Union for Conservation of Nature (IUCN) 2016, due to consequence of ongoing loss of their wetland habitats (Chutipong et al. 2019) and consequences of global climate change. Understanding the composition of excretions of semiochemicals that facilitate communication among fishing cats, as well as how they utilize chemical signals to navigate their environment and to search and interact with neighbouring potential fellow-mates and also for establishing territories, could provide critical insights for conservation efforts and initiatives. This paper aims to explore different behavioural connotations for pheromonal communication of the fishing cat, emphasizing on functional roles of specific semiochemical compounds associated with their marking modalities and social interactions. These molecules conceive information for coding by releaser and decoding by the recipient individuals about the reproductive status. The efficacy of these compounds to be remain stable when in Nature will investigate to extrapolate hypothetically the adaptive mechanism of fishing cat in wet-land ecology.

## Materials and methods

The possible pheromonal sources which were already been reported in other felids were taken into consideration for different experiments for FC. Fishing cats urinate normally in two modes, one upward with raising tail and in a violent jet, we designate this behavioural posture as “Marking Fluid (MF)” spray like that of other felid members (Das 2024); another is normal urination (UR). Samples from both these behavioural postures of FC were collected separately from Alipore zoological garden (Rescue centre of Zoo Authority), Kolkata, West Bengal, India (22.535913°N, 88.332053°E) and Garchumuk mini zoo and Rescue Centre, Howrah, West Bengal, India (22.348414°N, 88.075321°E) **(Online resource 1)** in the morning schedule (6am-8am), when resident animals were in their fully active state, with the assistance of known zoo-keepers. Samples were collected in a clean autoclaved beaker with the help of a clean sterilized glass-made Pasture pipette and were transferred into 10 ml air tight glass vials (Agilent, USA), crimped immediately with siliconized septum and transported to laboratory under ice for further analysis. Animals selected for the sample collections for experiments was mainly based on permission of zoo-authority. We were constrained by logistic issues, and other unavoidable factors for working with zoo-captive animals, which obviously restricted our choice. In every case, the samples were transported under ice to the laboratory and then preserved at -20°C for few days and then at -70°C for future analysis.

### For Identification of Volatile Organic Compounds (VOCs) from UR and MF of FC by Head-Space-Solid-Phase-Micro-Extraction–guided Gas Chromatography Mass Spectrometry (HS-SPME-GCMS)

A 1 cm 50/30 µm divinylbenzene/carboxen/polydimethylsiloxane (DVB/CAR/PDMS) solid phase microextraction (SPME) fiber (Supelco, U.S.A, stableflexTM, 24 Ga), which is suitable for collection of various VOCs from vapour phase for carbon number C3-20 and with MW 40-275, was chosen for the absorption of Head Space Volatiles (HSVs). In each case, 3 ml UR or MF was used for collection of HSVs by attaching the SPME fiber with the help of a manual assembly holder (Supelco, U.S.A.) positioned over the vials. Equilibration for adsorbing the vapor phase was maintained at room temperature for 1 hour. HSVs were then analysed using GC-MS after desorption to the injection port. Two GC-MS systems with two different columns were used for repeated confirmation: an Agilent 7890A (USA) equipped with a triple axis MS-5975C and a DB-WAX column (30 m × 0.25 mm × 0.25 µm); and a Thermo Fisher Trace 1300 (Thermo Scientific, Milan, Italy) connected to a triple quadrupole mass spectrometer (Thermo MS-TSQ 9000) and, with a wall-coated open tubular column (WCOT), TG-5 MS (30 m × 0.25 mm × 0.25 µm), a 10 m Dura guard capillary column. Samples were desorbed for 8 minutes at the injector port, which was maintained at 250°C. The flow rate of carrier gas Helium (purity >99.99%) was 1 mL/min. The column temperature started at 35°C for 2 minutes, ramped up to 210°C at a rate of 4°C/min, and was held at 210°C for 3 minutes in both cases. The MS source, quadrupole, and auxiliary heater temperatures were set at 230°C, 150°C, and 280°C, respectively. The electron energy was 70 eV (vacuum pressure 2.21 × 10^−5^ torr). Compound identification was achieved by comparing mass spectral data with the NIST MS search version 2.3 (2017) library and the NIST (2011) library. Identification was further confirmed by comparing mass fragmentation patterns with previous laboratory records and with some authentic standards [from Sigma (USA) and by Dr. Ehrenstorfer GmbH, (Germany)], whenever available and by calculating Linear Retention Index (LRI) in relation to n-alkanes of C8–C19.

### Estimation of less-volatile compounds from UR and MF of Fishing cat

10 µl of UR and MF samples were diluted with 1ml deionized water. Then, 250µl of 8% Ninhydrin (SRL, India) solution in ethanol was added and the mixture was kept in water bath at ±95°C for 15 min. The solution was turned into blue color forming Ruhemann’s complex (Ruhemann 1910). The micro-centrifuge tubes were cool down at room temperature and 250µl of absolute ethanol (Bengal Chemical, India) was added to the solution before measuring the optical density at 570 nm (Shimadzu, UV-Vis Spectrophotometer, UV-2600). Total free amines were estimated by developing standard curve against Glycine (Hi media, India).

Total proteins from the UR and MF were precipitated by adding chilled acetone (HPLC grade) to each sample separately in the ratio of 1:4, kept overnight at -20°C then centrifuged at 12000 rpm for 15 mins at 4°C (Eppendorf 5810R, USA). The precipitate was allowed to be air-dried and dissolved in 1 ml of 1M Tris-HCl buffer (pH 6.8). The protein was estimated spectrophotometrically at 595 nm against Bovine Serum Albumin in different dilution following Bradford assay (Bradford 1976).

Lipid was extracted from 5 ml of UR taken within a separating funnel, following modified Bligh and Dyer’s method (Bligh and Dyer 1959; Poddar-Sarkar 1996). Lower lipid-rich chloroform phase was subjected to High Performance Thin Layer Chromatography (HPTLC; CAMAG Linomat 5-160424) for separation of neutral lipid (NL) classes. Pre-coated silica plates (TLC plate; Merck, India - 10 × 10 cm or 10 × 20 cm Silica gel 60 F_254_) were used for spotting of extracted lipid and for development with mobile phase (Petroleum ether: Diethyl ether: Acetic acid in the ratio of 70:30:1.5 or 80:20:1.5 or 90:10:1.5, whenever required for better separation of bands). After development, plates were sprayed with 0.2 % (w/v) solution of 2,7-dichlorofluorescein (DCF) in methanol and visualized under 366nm by the detector (CAMAG TLC Scanner 3-180418) using scanning speed 20mm/s, Lamp – Mercury (Hg), Slit dimension - 6.00 mm x 0.30 mm). Identification was done by comparing Rf values with authentic compounds (Sigma, USA) such as Diacylglyceride (DAG/DG, as Dipalmitin), Cholesterol (CHOL, as Free sterol), Long chain alcohol (LCA, as Eicosanol), Free fatty acid (FFA, as Decanoic acid) Triacylglyceride (TAG/TG, as Tripalmitin), Wax-ester (WE, as Stearyl arachidate), Sterol ester (SE, as Cholesteryl palmitate) developed in the same manner. Quantification from samples of FC was done by developing respective standard curve with specific concentration of authentic compounds with the help of WinCATS Planner Chromatography Manager Software and expressed in µg/ml.

Another aliquot of lower lipid-rich chloroform phase was taken for profiling of Fatty acids (FAs) from the lipid part of UR and MF (Chalvardjian and Still 1964; Poddar-Sarkar 1996). A portion of the extracted chloroform from UR and MF was concentrated with a stream of nitrogen to 0.5 ml. A mixture of methanol, benzene, and concentrated HSO_4_ (4.3:0.5:0.8) was added to the chloroform phase for acid hydrolysis (Christie 1993; Poddar-Sarkar 1996). The solution was heated at 90°-95°C in a water bath for 8 hours and then left at room temperature overnight. The fatty acid methyl esters (FAMEs) were extracted from sap fraction with n-hexane (E. Merck, India, HPLC grade) and dried over anhydrous sodium sulphate. The volume of the n-Hexane was reduced to the required concentration with a stream of nitrogen before injecting into GC-MS column for analysis of FAMEs. 1 µl of hexane containing FAMEs was injected into a HP5-MS capillary column (30 m × 0.25 mm × 0.25 μm; Agilent), with a 10 m Duraguard, connected to an Agilent 7890A coupled with a triple-axis mass detector (Agilent MS-5975C) for analysis. The temperature program started at 70°C, held for 1 minute, then ramped up at 4°C/min to 260°C, by keeping He flow at 1ml/min. and held at this final temperature for 3 minutes. FAMEs were identified by comparing their mass fragmentation patterns to those in the NIST (2011) library, and confirmation was achieved by calculating relative retention times (RRt) against 37 FAME and PUFA authentic standard mixtures (Supelco, Lot No: LB80556 & LB77207, USA).

### Exploration on behavioral strategy

After identification of HSVs by GC-MS through SPME from UR and MF, we tried to explore the strategic modalities which has been evolved in fishing cats, impacted by semi aquatic-ecoenvironment for adaptation and which help them for survival in swapmy-marshy habitats. During our close observation we found that FCs very frequently urinate in waterbodies of the enclosures or within drinking water-pots kept inside the enclosure for them. Thus, we searched for logical resoning behind this fact of natural phenomenon and attempted to simulate by formulating two experiments. It was also noted from GCMS data that UR & MF of FC contain high amount of hydroxy-functional group. From this prediction, for the first experiment we collected two available samples from the water-pot to investigate what types of compounds are coming out in HSVs from the water content (ignoring the exact concentration of the compounds). We took water-content from pot into an air-tight, teflon-coated glass vial, crimpped instantly; and SPME fibre was attached to draw HSVs for subsequent analysis by GC-MS following above procedure. In next step, as we already obtained from GC-MS data that the appearance of marker-pheromone of FC, ‘Tom cat compound’ 3-Mercapto-3-methyl-butanol (3MMB) and other surfur derivatives is a time-dependent process (Das et al. 2023b) which are produced by break-down of precursor amino acid Felinine, either by physical factors or by bacterial degration (Das et al. 2023b), we formulate second experiment. Felinine (2-Amino-7-hydroxy-5,5-dimethyl-4-thiaheptanoic acid) is also identified as the precursor of pheromone in domestic cats, bob-cats urine and Eurasian lynx (Miyazaki et al. 2006a). For this experiment, we use synthetic 3MMB which is a sulfur containing alcohol and with high affinity towards water having low hydrophobicity (logp= 0.93). We scheduled experimental set-up for 21 days span at laboratory atmosphere to check the retention ability of 3MMB in water, pretending the possible changes when in nartural environment. It is to be noted that bioassay by applying directly to animals for investigations was not permissible by the guidelines of zoo authority. For this purpose, we added 1µl of 3MMB standard (Sigma, USA) in 10 ml of distilled water and transfer into a wide-mouth, open-cap glass-vial and kept at laboratory environment for natural transformation. Then, we performed repeated head-space-GC-MS in 7 days interval upto 21 days by absorbing through SPME following the same procedure and subsequently recorded the relative Total Ion Count (TIC) from the chromatogram.

## Results

### VOCs of Urine & Marking Fluid of Fishing cats

GC-MS analysis of UR of FC revealed the presence of various functional groups, including alcohols, ketones, aldehydes, and sulfur- and nitrogen-containing compounds in headspace sampling from both sexes. Ketones were the most prevalent group (male: 37.83 ± 21.95%; female: 43.97 ± 19.07%), followed by aldehydes (male: 32.17 ± 25.08%; female: 27.42 ± 7.15%) and sulfur-containing compounds (male: 17.65 ± 13.43%; female: 1.46 ± 0.22%) **(Fig. 1)**. A total of 42 VOCs from UR of male fishing cat (from 11 sampling) were identified (**Table 1**), among which 11 compounds including 3-Buten-1-ol, 3-methyl-; 2-Buten-1-ol, 3-methyl; Phenol; Benzaldehyde; 2-Pentanone; 4-Heptanone; Acetophenone; 3-Mercapto-3-methylbutyl formate (3MMBF); 3-Mercapto-3-methyl-butanol (3MMB); 3-Methyl-3-methylthio-1-butanol (3MMBT); and 3-Methyl-3-(2-methyldisulfanyl)-1-butanol (3M3MDSB) were detected in most of the UR samples. Among these, Benzaldehyde (29.68 ± 24.96 %), 4-Heptanone (17.61 ± 13.39 %), 3MMB (10.35 ± 11.66%), Acetophenone (5.58 ± 2.05 %) and 2-Pentanone (6.74 ± 7.33 %) exhibited notably high mean abundances in the urinary headspace VOCs.

**Table 1.** List of HSVs identified from chromatogram of UR and MF of FC. Top five characteristic m/z and linear retention indices of identified compounds are given as reference. [Presence in male and female samples are represented with m and f (superscript) respectively].

| Head space volatiles (HSVs) |  | Occurrence |  |  |  | Top five mass fragmentation pattern (m/z) | Linear retention index (LRI) |
| --- | --- | --- | --- | --- | --- | --- | --- |
| Volatile classes | Compounds | UR |  | MF |  |  |  |
|  |  | DB-WAX | TG-5 | DB-WAX | TG-5 |  |  |
| Alcohols | 3-Buten-1-ol, 3-methyl-<sup>m,f</sup> | ✓ | ✓ | ✓ | ✓ | 41(BP), 56, 68, 86, 67 | 727 (TG-5) |
|  | 2-Buten-1-ol, 3-methyl-<sup>m,f</sup> | ✓ | ✓ | ✓ | ✓ | 71(BP), 41, 43, 86, 53 | 773 (TG-5) |
|  | 1-Hexanol<sup>m</sup> | ✓ |  | ✓ |  | 56(BP), 43, 41, 55, 42 | 1359 (DB-wax) |
|  | 1-Heptanol<sup>m</sup> | ✓ |  | ✓ |  | 70(BP), 56, 43, 41, 55 |  |
|  | 2-Ethyl-1-hexanol<sup>m,f</sup> | ✓ |  | ✓ |  | 57(BP), 41, 43, 70, 55 | 1496 (DB-wax) |
|  | 1-Octanol<sup>m,f</sup> | ✓ |  | ✓ |  | 56(BP), 55, 41, 43, 70 | 1565 (DB-wax) |
|  | Benzenemethanol, alpha.-methyl-<sup>m</sup> | ✓ | ✓ | ✓ |  | 105(BP), 77, 78, 89, 107 |  |
|  | Phenol<sup>m,f</sup> | ✓ | ✓ | ✓ |  | 94(BP), 66, 65, 55, 95 | 980 (TG-5) |
|  | p-Cresol<sup>m,f</sup> | ✓ |  | ✓ |  | 107(BP), 108, 77, 79, 109 | 1060 (TG-5) |
| Aldehydes | Hexanal<sup>m,f</sup> | ✓ | ✓ |  |  | 44(BP), 56, 41, 57, 43 | 1186 (DB-wax); 799 (TG-5) |
|  | Heptanal <sup>m</sup> | ✓ | ✓ |  |  | 70(BP), 55, 44, 43, 57 | 1182 (DB-wax); 904 (TG-5) |
|  | 2-Butenal, 3-methyl- <sup>m</sup> | ✓ | ✓ | ✓ |  | 84(BP), 83, 55, 41, 53 | 791 (TG-5) |
|  | Nonanal <sup>m,f</sup> | ✓ | ✓ | ✓ | ✓ | 57(BP), 56, 70, 98, 41 | 1391 (DB-wax); 1107 (TG-5) |
|  | Benzaldehyde <sup>m,f</sup> | ✓ | ✓ | ✓ | ✓ | 77(BP), 105, 106, 44, 51 | 1516 (DB-wax); 963 (TG-5) |
| Ketones | 2-Hexanone, 3-methyl- <sup>m</sup> | ✓ |  |  |  | 43(BP), 72, 71, 55, 70 |  |
|  | 2-Pentanone <sup>m,f</sup> | ✓ | ✓ | ✓ | ✓ | 43(BP), 86, 71, 58, 41 |  |
|  | 2-Pentanone, 3-ethyl- <sup>m</sup> | ✓ | ✓ | ✓ | ✓ | 43(BP), 41, 57, 56, 72 | 861 (TG-5) |
|  | 4-Heptanone <sup>m,f</sup> | ✓ | ✓ | ✓ | ✓ | 43(BP), 71, 114, 41, 58 | 1118 (DB-Wax); 837 (TG-5) |
|  | 2-Heptanone <sup>m</sup> | ✓ | ✓ |  | ✓ | 43(BP), 58, 71, 59, 114 | 855 (TG-5) |
|  | 3-Octanone <sup>m</sup> | ✓ | ✓ |  |  | 43(BP), 57, 72, 71, 99 | 986 (TG-5) |
|  | 2-Octanone <sup>m,f</sup> | ✓ |  | ✓ |  | 43(BP), 58, 41, 71, 81 | 1284 (DB-Wax); 955 (TG-5) |
|  | 2-Hexanone, 3,4-dimethyl- <sup>m,f</sup> | ✓ | ✓ | ✓ |  | 43(BP), 72, 41, 85, 57 |  |
|  | 4-Nonanone <sup>m,f</sup> | ✓ | ✓ |  | ✓ | 43(BP), 71, 58, 99, 86 |  |
|  | 2-Nonanone <sup>m,f</sup> | ✓ | ✓ | ✓ | ✓ | 43(BP), 58, 41, 57, 71 | 1387 (DB-wax); 1093 (TG-5) |
|  | 2-Decanone <sup>m</sup> | ✓ |  |  |  | 58(BP), 43, 59, 71, 85 |  |
|  | 2-Undecanone <sup>m,f</sup> | ✓ | ✓ |  |  | 58(BP), 43, 71, 59, 81 | 1296 (TG-5) |
|  | Acetophenone <sup>m,f</sup> | ✓ | ✓ | ✓ | ✓ | 105(BP), 77, 120, 43, 51 | 1645 (DB-wax); 1072 (TG-5) |
| Sulfur containing compounds | 3-Methyl-3-butene-1-thiol <sup>m</sup> |  | ✓ |  | ✓ | 102(BP), 41, 44, 56, 59 |  |
|  | Dimethyl disulfide <sup>m,f</sup> | ✓ |  |  |  | 94(BP), 79, 45, 46, 47 |  |
|  | Thiophene, 2-methyl- <sup>m</sup> | ✓ | ✓ |  |  | 97(BP), 98, 44, 43, 45 | 748 (TG-5) |
|  | Thiophene-2-propyl- <sup>m</sup> | ✓ |  |  |  | 97(BP), 126, 98, 99, 45 |  |
|  | Thiophene, 2-methoxy-5-methyl- <sup>m,f</sup> | ✓ |  | ✓ |  | 128(BP), 113, 85, 69, 59 |  |
|  | 3-Mercapto-3-methylbutyl formate <sup>m</sup> | ✓ | ✓ | ✓ | ✓ | 69(BP), 43, 103, 41, 75 | 1036 (TG-5) |
|  | 3-Mercapto-3-methyl-butanol <sup>m</sup> | ✓ | ✓ |  | ✓ | 41(BP), 69, 86, 71, 120 | 972 (TG-5) |
|  | 3-Methyl-3-methylthio-1-butanol <sup>m</sup> | ✓ | ✓ | ✓ | ✓ | 69(BP), 41, 134, 93, 71 | 1160 (TG-5) |
|  | 3-Methyl-3-(2-methyldisulfanyl)-1-butanol <sup>m,f</sup> | ✓ | ✓ | ✓ | ✓ | 69(BP), 41, 87, 80, 148 | 1334 (TG-5) |
| Nitrogen containing Compounds |  |  |  |  |  |  |  |
|  | Pyrazine, 2,5-dimethyl- <sup>m,f</sup> | ✓ |  |  |  | 42(BP), 108, 81, 43, 40 |  |
|  | Pyridine, 2,4,6-trimethyl- <sup>m,f</sup> | ✓ |  | ✓ |  | 121(BP), 79, 106, 120, 77 |  |
|  | Heptanonitrile <sup>m,f</sup> | ✓ |  |  |  | 41(BP), 82, 83, 55, 54 |  |
|  | Indole <sup>m,f</sup> | ✓ | ✓ | ✓ | ✓ | 117(BP), 90, 89, 116, 118 | 1301 (TG-5) |
| Others |  |  |  |  |  |  |  |
|  | Acetic acid <sup>m</sup> | ✓ |  |  |  | 43(BP), 60, 45, 57, 42 | 1427 (DB-Wax) |
|  | Azulene <sup>f</sup> | ✓ | ✓ | ✓ |  | 146(BP), 148, 111, 113, 75 | 1729 (DB-Wax); 1330 (TG-5) |

**Fig. 1.**
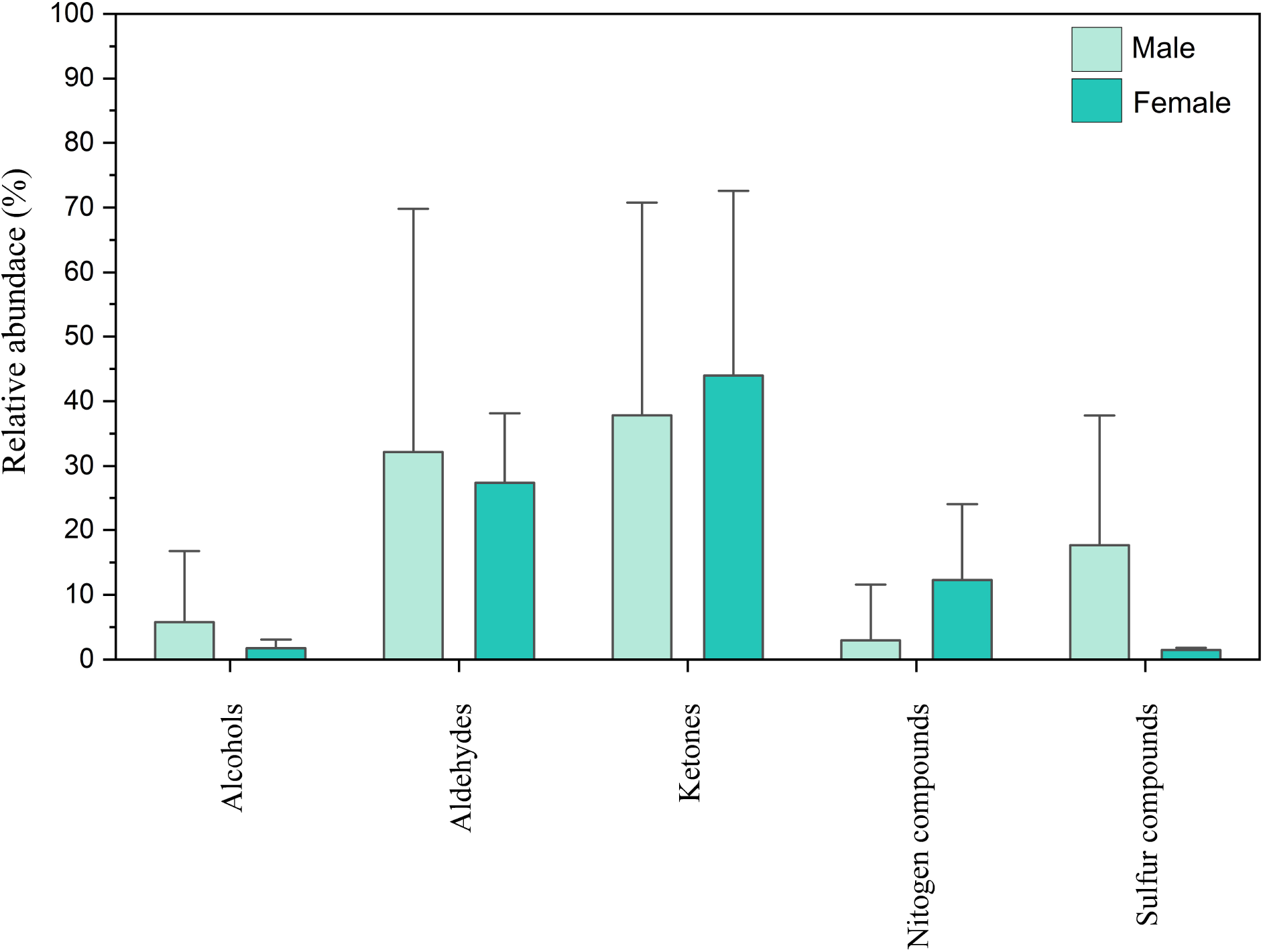
Box plot showing the cumulative abundance of each functional groups detected from urine of fishing cat of male and females.

In two female UR samples, a total of 25 VOCs were detected among which Benzaldehyde (24.95 ± 6.67%), 4-Heptanone (17.34 ± 2.32%), Acetophenone (13.55 ± 13.03%), 2-Octanone (7.22 ± 4.05%) were most dominant HSVs. It is interesting to note that 3MMB was detected from all the male UR samples, but not from female UR samples **(Fig 2)**. Although the absence of 3MMB in female samples cannot be concluded at this stage due to small sample size (two samplings), because of our limited access to collection from female FC as they are isolated for breeding programme by zoo. Miyazaki *et al*., (2006) & (2008) has given a hint for occurrence specificity of this compound only in male domestic cat urine. Similar to 3MMB, other sulfur-containing VOCs like 3MMBF (from 9 sample out of total 11 samples) and 3MMBT (11 out of 11 samples) were also detected only in male UR samples **(Online resource 2)**. On the other hand, sulfur-containing compound 3M3MDSB was detected in both male and female UR samples.

**Fig. 2.**
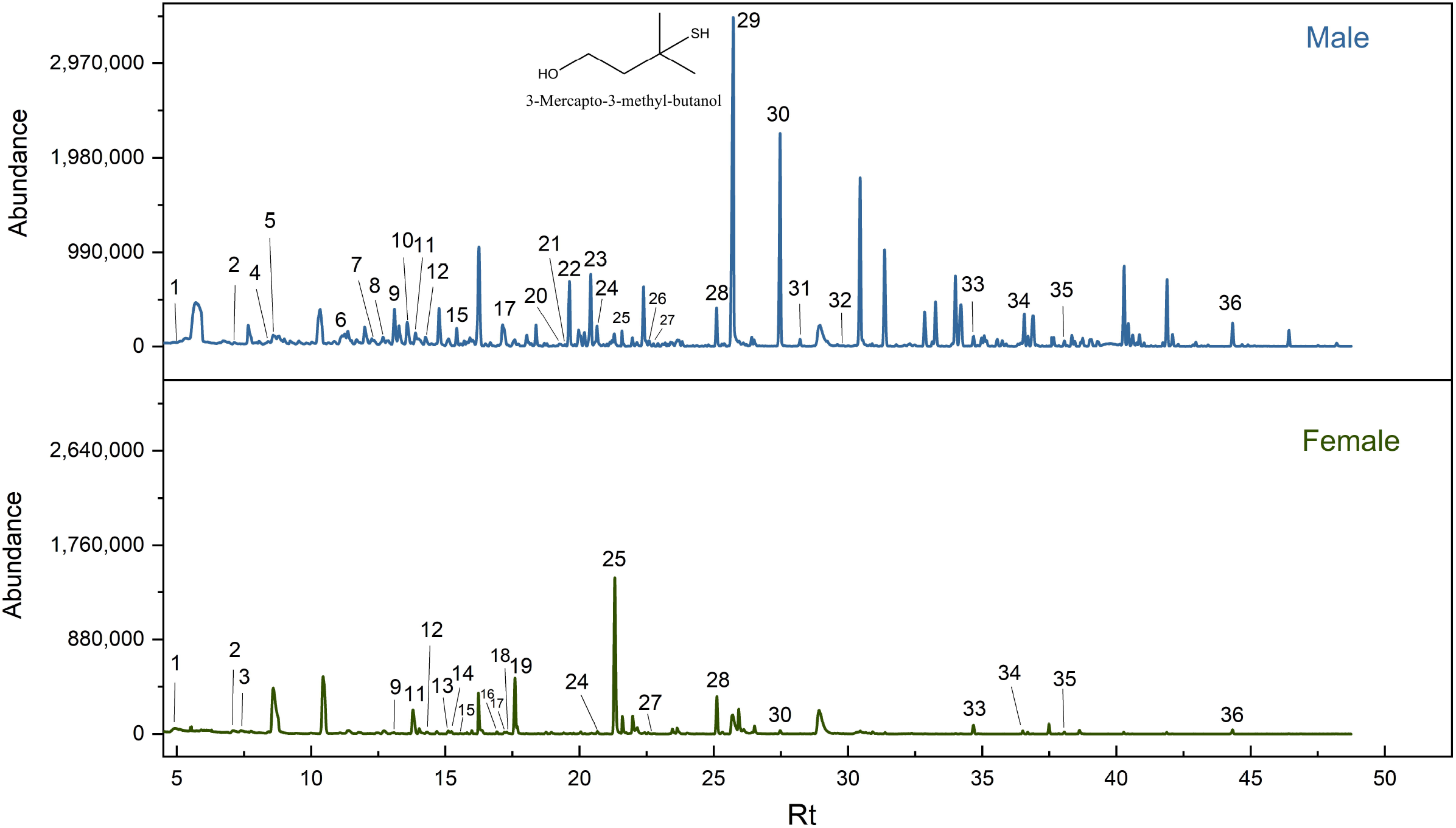
Gas chromatograms obtained using a DB-Wax column showing a representative HSV profile from the urine of male and female fishing cats (X axis=Retention time in min, Rt; Y = relative abundance). 1. 2-Pentanone; 2. Dimethyl disulfide; 3. Hexanal; 4. Thiophene, 2-methyl-; 5. 4-Heptanone; 6. 2-Butenal, 3-methyl-; 7. Thiophene-2-propyl-; 8. 3-Octanone; 9. 3-Buten-1-ol, 3-methyl-; 10. 2,4-Dithiapentane; 11. 2-Octanone; 12. 2-Hexanone,3,4-dimethyl-; 13. 4-Nonanone; 14. Pyrazine, 2,5-dimethyl-; 15. 2-Buten-1-ol, 3-methyl-; 16. Pyridine, 2,4,6-trimethyl-; 17. 2-Nonanone; 18. Nonanal; 19. Heptanonitrile; 20. Thiophene-2-pentyl-; 21. Acetic acid; 22. 1-Heptanol; 23. 2-Decanone; 24. 2-Ethyl-1-hexanol; 25. Benzaldehyde; 26. 3-Mercapto-3-methylbutyl-formate; 27. 1-Octanol; 28. Acetophenone; 29. 3-Mercapto-3-methyl-butanol; 30. Azulene; 31. 3-Methyl-3-methylthio-1-butanol; 32. Benzenemethanol, .alpha.-methyl-; 33. Phenol; 34. p-Cresol; 35. 3-Methyl-3-(2-methyldisulfanyl)-1-butanol; 36. Indole.

Analysis of HSVs from male MF samples revealed presence of 30 VOCs (from 5 samples) **(Table 1)**, where Benzaldehyde (57.59 ± 4.46 %) was the most dominant VOC **(Fig. 3; Online resource 3)**. All the HSVs that were detected in MF samples were also detected in UR samples. Comparative study of relative abundance of VOCs between male UR and MF by Mann-Whitney test revealed that relative abundance of Benzaldehyde (Mann-Whitney U= 6; z= 2.3791; p=0.017) and 3MMBT (U= 0; z= 3.061; p=0.002) is significantly high in MF than UR. There are no differences in the abundance of 4-Heptanone, 3MMB in male UR and MF samples.

**Fig. 3.**
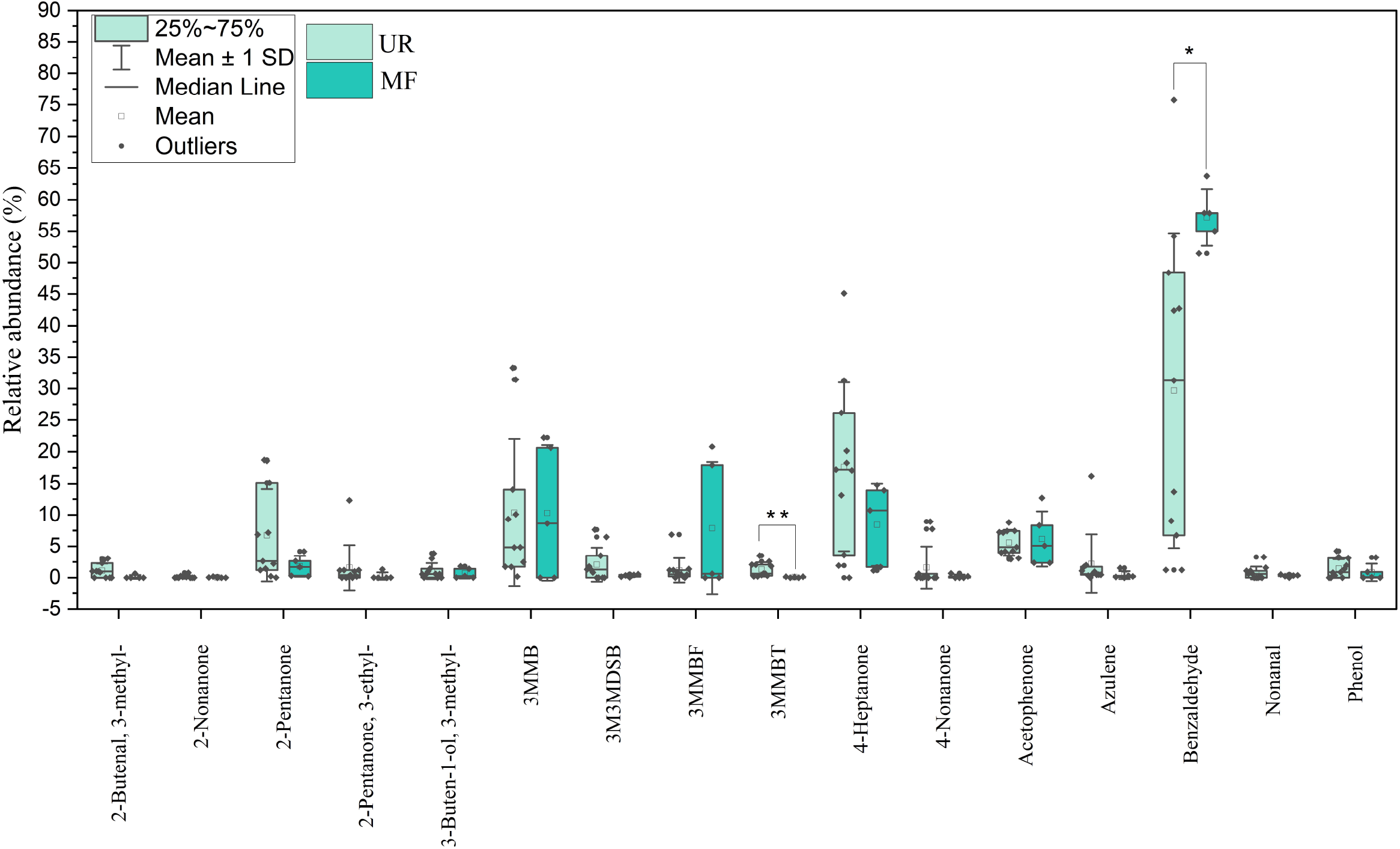
Box plot showing relative abundance of 16 frequently (detected in more than 50% of the sample run) identified compounds in HSVs of UR and MF of male FC. Significant differences were represented by asterisk; (p<0.05=*; p<0.01=**)

### Analysis of less volatile compounds in Urine & Marking Fluid of Fishing cats

The UR of FC males contained higher free amines (3.33±1 mg/ml) but lower soluble proteins (0.15± 0.2 mg/ml) in compared to MF samples (free amines: 2.07±0.3 mg/ml; soluble proteins: 0.32± 0.2 mg/ml). The amount of total lipid in UR and MF of males were 1.415 ± 0.007 mg/ml and 1.995 ± 0.062 mg/ml respectively. The concentrations of NL classes estimated from HPTLC analysis, calculated against standards from UR of FC were Diacylglycerol - 20.8±1.3µg/ml, Cholesterol-7.8±4µg/ml, Free Fatty Acids - 23±9.4µg/ml and Triglyceride - 78.7±10.8µg/ml **(Fig. 4)**.

**Fig. 4.**
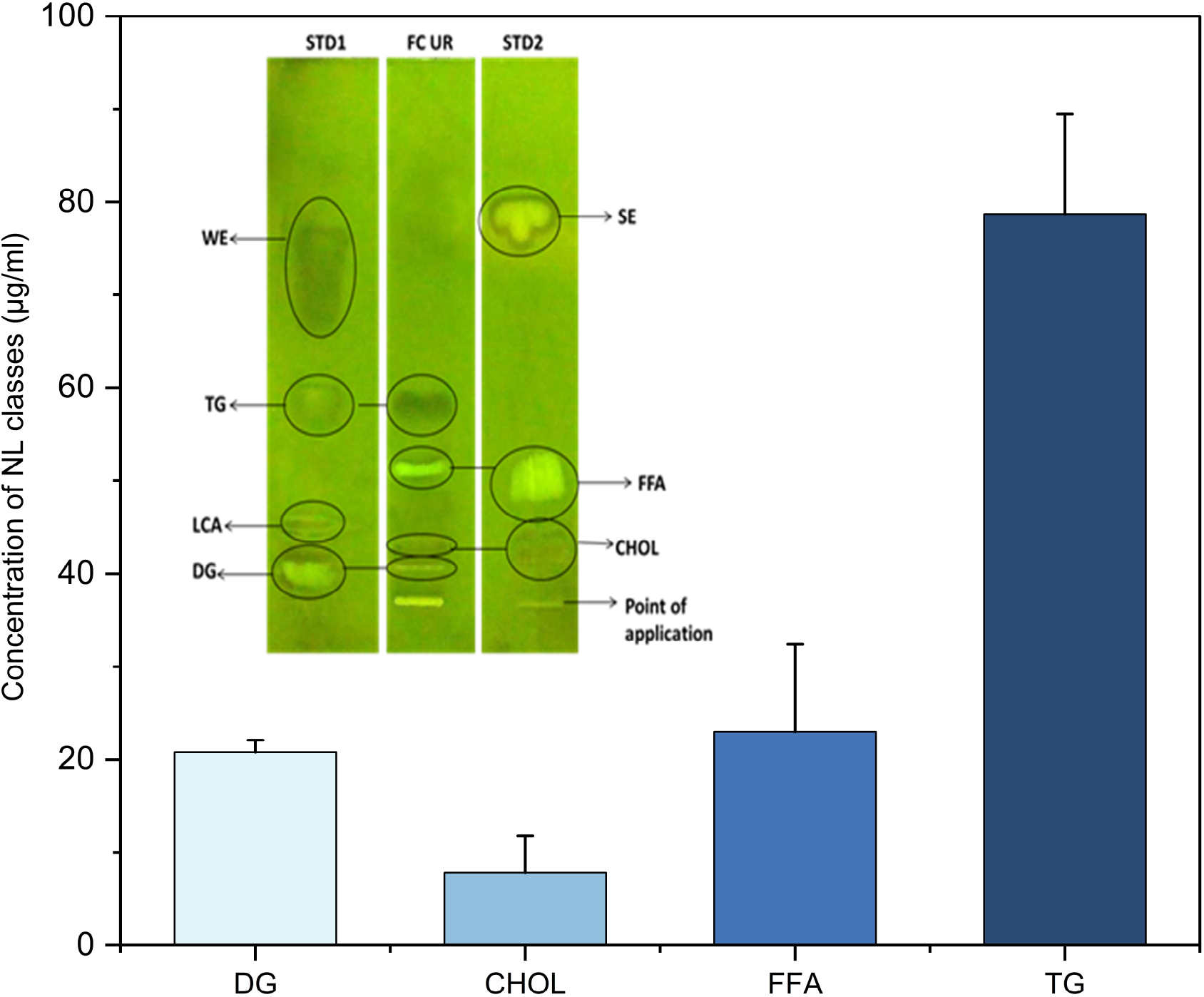
Histogram showing concentrations of Diglyceride, Cholesterol, Free fatty Acids and Triglyceride. HPTLC plates (inset) developed from lipid extract of UR of FC showing bands of neutral lipid classes (Standard mixture of DG-Diglyceride, LCA-Long chain alcohol, TG-Triglyceride, WE-Wax ester in STD-1 lane, UR-FC-lipid in lane 2 and Standard mixture of CHOL-Cholesterol, FFA-Free fatty Acids, SE-Sterol ester in STD 3 lane).

From the lipid portion of the male UR and MF, saturated (SFA), monounsaturated (MUFA), and polyunsaturated fatty acids (PUFA) were detected. SFAs ranging from C10-C24 were identified in UR and MF **(Online resource 4)**. Hexadecanoic acid (C16:0) is the most abundant in UR (30.84 ± 6.57%) and MF (33.61 ± 3.81%) followed of Octadecanoic acid (C18:0 UR: 12.51 ± 3.3%; MF: 13.59 ± 2.52%) in case of SFAs. The most abundant UFA is 11-Octadecenoic acid [C18:1(Δ11) UR: 20.56 ± 6.25%; MF: 10.83 ± 2.27%]. Besides, the other MUFAs identified from UR and MF are 9-Hexadecenoic acid, 9-Octadecenoic acid, 11-Eicosenoic acid, and 13-Docosenoic acid. Three PUFAs that were identified from both UR and MF of fishing cat are 5,8,11-Eicosatrienoic acid; 9,12-Octadecadienoic acid; and 11,14 Eicosadienoic acid **(Fig. 5)**.

**Fig. 5.**
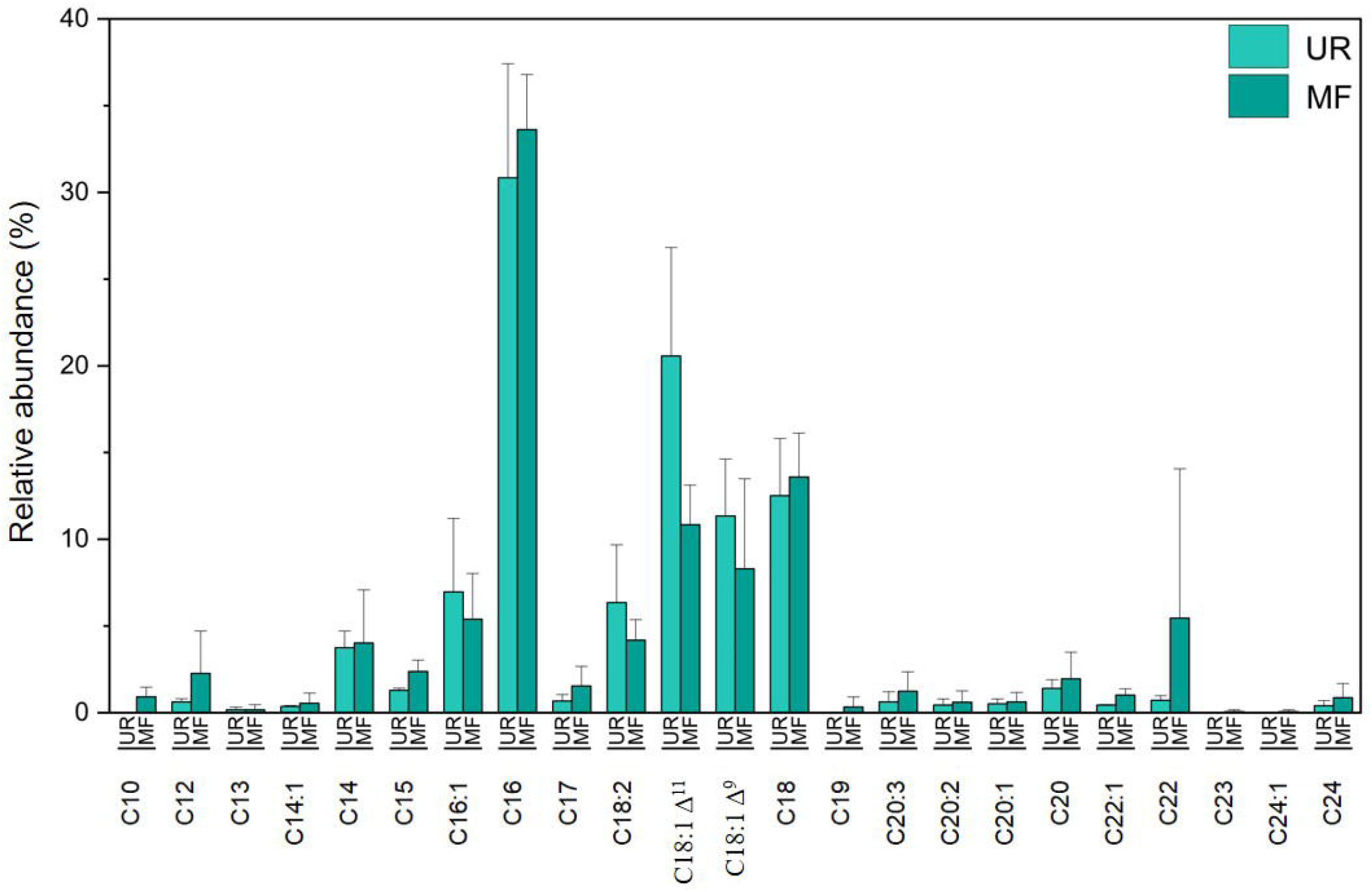
Relative abundance of identified fatty acids from UR and MF of male FC. In the bar diagram, each fatty acid is represented with their respective carbon numbers C10-C24 in x-axis.

### Exploration on behavioral strategy

The GC-MS chromatogram of first experiment i.e head space drawn from water content, detected 4 and 7 VOCs in two samples respectively. These were 3MMB from two collections (i.e 2/2), 2-Pentanone (from 1 sample), 4-Heptanone (from 2/2), 2,4-Dithiapentane (from 1/2), Hexanal (from 1/2), 3MMBF (from 1/2) and Azulene (from 1/2). Surprisingly, among all the detected HSVs, the relative abundance of 3MMB is exceptionally high (70.44 ± 2.69 %) **(Fig 6)**. The retention of 3MMB is possibly due to its high affinity towards water. **(Fig. 6)**. The retention of 3MMB is possibly due to its high affinity towards water. In a follow-up experiment aimed at assessing the retention ability of synthetic 3MMB in water over a 21-day period, results demonstrated that 3MMB remained substantially retained, with a reasonable TIC signal persisting at 28.5% of the initial signal by the end of the study. **(Fig. 7)**.

**Fig. 6.**
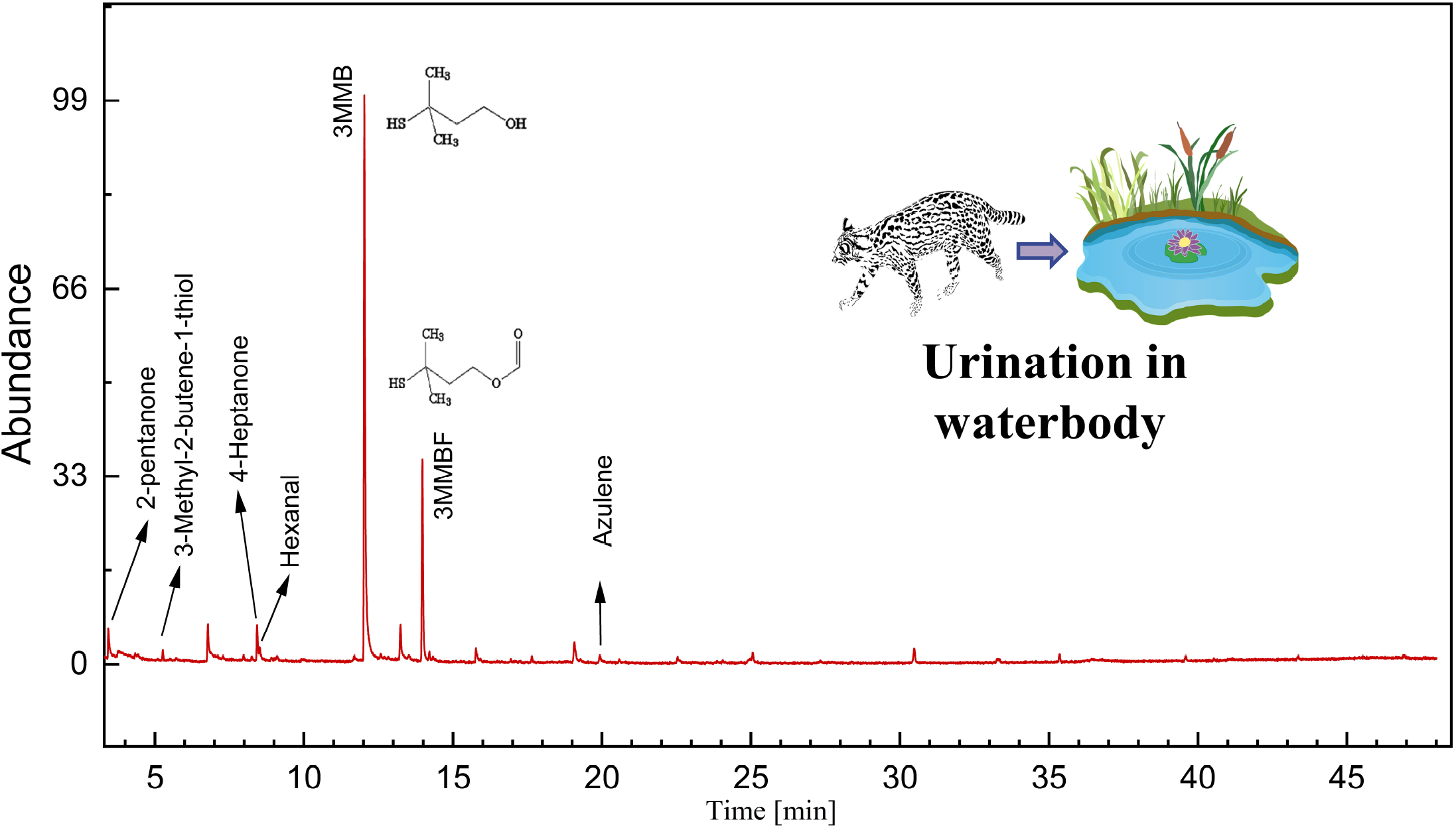
GC-MS chromatogram illustrating HSV profiles from water-content containing UR of fishing cat.

**Fig. 7.**
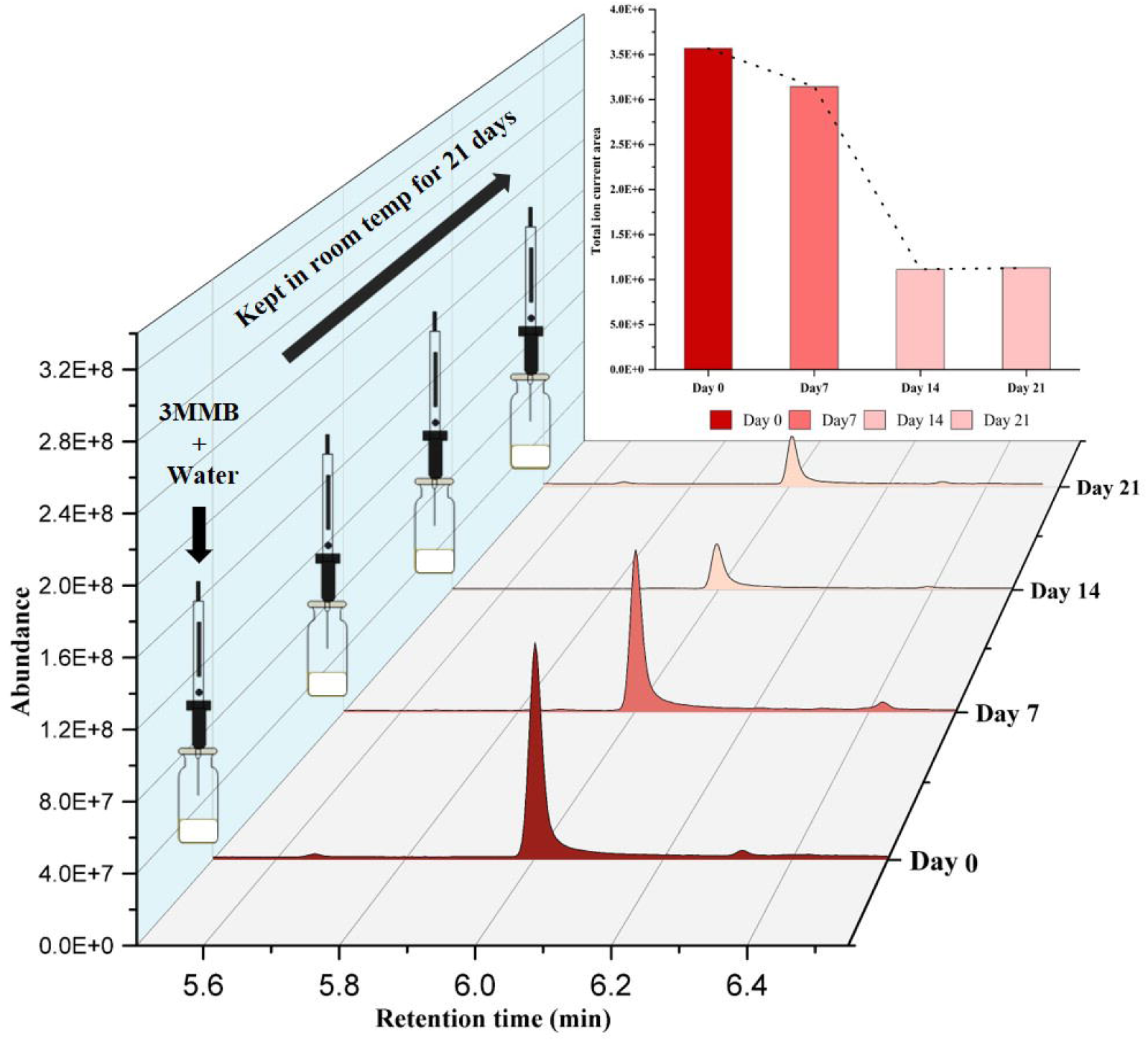
Intensity and TIC area (inset) of 3MMB in GC chromatograms obtained from water-urine content of fishing cat analysed on 0^th^, 7^th^, 14^th^ and 21^st^ days.

## Discussion

These identified VOCs from UR and MF of fishing cats, synthesized by metabolic routes contribute significantly in chemical communication system of this tropical cat member, adapted in semi-aquatic environment. The similarity in the chemical composition of UR and MF suggests that the compounds in both sources are derived from a common *in vivo* biochemical pathway and might be excreted through a common excretory duct. Several compounds present in UR and MF of FC including 3MMB, Benzaldehyde, Indole, Phenol, 3-Methyl-3-buten-1-ol, and 2-Methyl-3-buten-1-ol, which were also identified from other felids and are supposed to play the significant role in the chemical communication (Miyazaki et al. 2006b, 2008; Soso and Koziel 2016, 2017; Das et al. 2023b; Das 2024). The sulfur-containing felinine breakdown volatile, 3MMB, is likely a sex-specific compound in fishing cats, and its biosynthesis might be regulated by male-specific hormone testosterone (Hendriks et al. 1995; Miyazaki et al. 2006a). It has been reported by many scientists that the detection of 3MMB is largely restricted to small felids, including domestic cats (Miyazaki et al. 2006a, 2008), leopard cats (Ichizawa et al. 2023) and lynxs (Mattina et al. 1991), where as in contrast, in large felids, such as in cheetah, tiger, jaguar, puma, Indian leopard, clouded leopard, lion, and snow leopard, 3MMB and other sulfur compounds associated with felinine have not been occurred (Poddar-Sarkar and Brahmachary 1997; Burger et al. 2006; McLean et al. 2007; Ghosh 2017; Das et al. 2019, 2023a). Thus, the detection of 3MMB, as a representative of breakdown product of felinine, in feline members may provide an important clue for tracing the evolutionary history of the global felid lineage; although, this specificity of occurrence of felinine as a chemical marker in small cats, but not in the big cat lineage, remains controversial.

As 3MMB is an important chemical compound for intraspecific and interspecific communication in felids, its sustainability in environment is essential for minimizing the necessity of scent-marking, thus for making energetically favourable for the concerned organism. As previously mentioned, field observation revealed that fishing cats frequently urinate in water, and our experiments confirm the high retention of 3MMB in water even after 21 days of application **(Fig. 6-7)**. Therefore, urination in water can be justified as an efficient strategy for prolonged presence of scent marks in marshy environments. It can be conceptualised from the fact that due to the slow release of this alcohol-containing VOC from water and for longer environmental sustainability, this procedure was favoured by natural selection and might have been settled evolutionarily, enabling fishing cats to economise metabolic energy for reducing frequency of scent marking for the conspecifics but to be coherent to be adopted in semi-aquatic environments. We conclude that it could be an excellent strategy for FC to mark on waterbodies to retain its specific odour in aquatic environments for a longer duration.

In contrast, the presence of less-volatile compounds, such as triglycerides, diglycerides, sterols, free fatty acids, and ensembles of long-chain fatty acids in the urine of fishing cats may extend the durability of chemical signals in nature or act as ‘fixatives’ for highly volatile molecules **(Fig. 4-5)**. These compounds likely bind with the low molecular weight, highly volatile molecules, stabilizing them in the terrestrial environment also and thereby enhancing the nature of complexity by degradation, which might be creating the purpose of imprinting ‘uniqueness’ for distinguishing self from non-self or for the aid of ‘overmarking’, which is the fundamental characteristic of pheromonal information.

In conclusion, unravelling the fact of chemical biochemical fingerprinting by Urine and Marking fluid of fishing cats opens up an intricate mechanism of chemical signalling, likely shaped by the challenges of their semi-aquatic lifestyle. The retention of 3MMB in water, along with the high abundance of polar compounds points out a highly effective strategy for scent marking in watery environments. Exploratory research with bioassays and behavioural studies may elucidate further, the role of these compounds in social interactions, territoriality and mating behaviour of fishing cats. These findings could be projected in broader perspectives of felid chemical communication system and may build up the path for strategic planning for conservation of this vulnerable species in future.

## Acknowledgments

Author SD [Award Number-08/155(0080)/2019-EMR-I)], SM [Award Number 08/155(0078)/2019-EMR-I] and PD [Award Number-09/028(1098)/2019-EMR-1] are grateful to Council of Scientific and Industrial Research (CSIR), Government of India for providing their fellowships during this work. We would also like to acknowledge the Department of Science and Technology, Government of India for proving GCMS facilities in Department of Botany, University of Calcutta [SR/FST/LSI-459/2010 dated 10.03.2011] and to the Department of Instrumentation and Electronics Engineering, Jadavpur University, [SR/FIST/ETI-424/2016 Date: 16.12.2016]. We are grateful to the Principal Chief Conservator of Forest (Wildlife) and Member Secretary, Zoo Authority, Government of West Bengal and the Director of Alipore zoo, Kolkata, India for their kind help and assistance.

## Statements and Declarations

### Conflict of interest

All authors have seen and approved the manuscript, and it has been not submitted to or published elsewhere. The authors have no relevant fnancial or non-fnancial interests to disclose. The authors declare that they have no conflict of interest.

## CRediT authorship contribution statement

Conceptualization: [Mousumi Poddar Sarkar]; Methodology: [Subhadeep Das and Payel Das]; Formal analysis and investigation: [Subhadeep Das and Sourav Manna], Writing - original draft preparation: [Subhadeep Das]; Writing-review and editing: [Sourav Manna and Mousumi Poddar Sarkar]; Funding acquisition: [Contingency of CSIR and self-funding], Resources: [Zoo animals of Zoo authority & Dept. of Forest, Govt.of West Bengal]; Supervision: [Mousumi Poddar Sarkar]

## Notes

### Competing Interest Statement

The authors have declared no competing interest.

