## Supplemental for "Functional insights on chemical communication of fishing cats: a strategic mechanism for adaptation in semi-aquatic habitat"

**Online resource 1: Details of the samples collected from Fishing cats during the whole span of study from 2017- 2022 in different field session (DOC- date of collection).**

| Samples from FC | DOC of UR | DOC of MF |
|---|---|---|
| **FCM1 (male) -AZG** | 09.05.2017 | 09.05.2017 |
| | 18.07.2018 | |
| **FCM2 (male)-AZG** | 12.05.2017 | 17.05.2017 |
| | 17.05.2017 | 21.08.2017 |
| **FCF1 (female)-AZG** | 12.04.2018 | |
| **FCM3 (male)-GMZ** | 26.02.2022 | |
| | 15.03.2022 | |
| **FCM4 (male)-GMZ** | 26.02.2022 | 26.02.2022 |

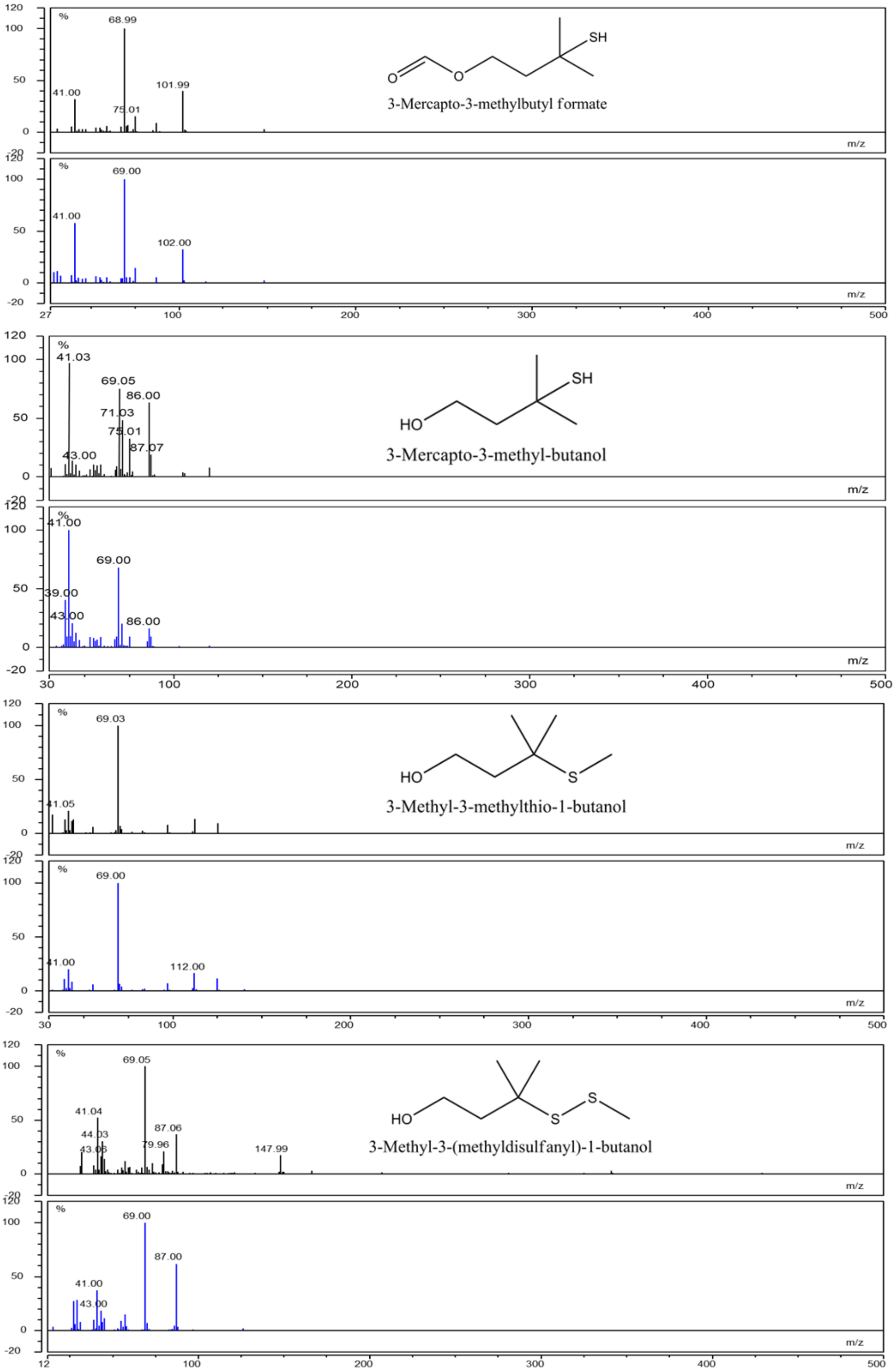


**Online resource 2: Mass Fragmentation pattern of felinine derivative compounds (black- compound mass pattern; blue-NIST library mass pattern)**

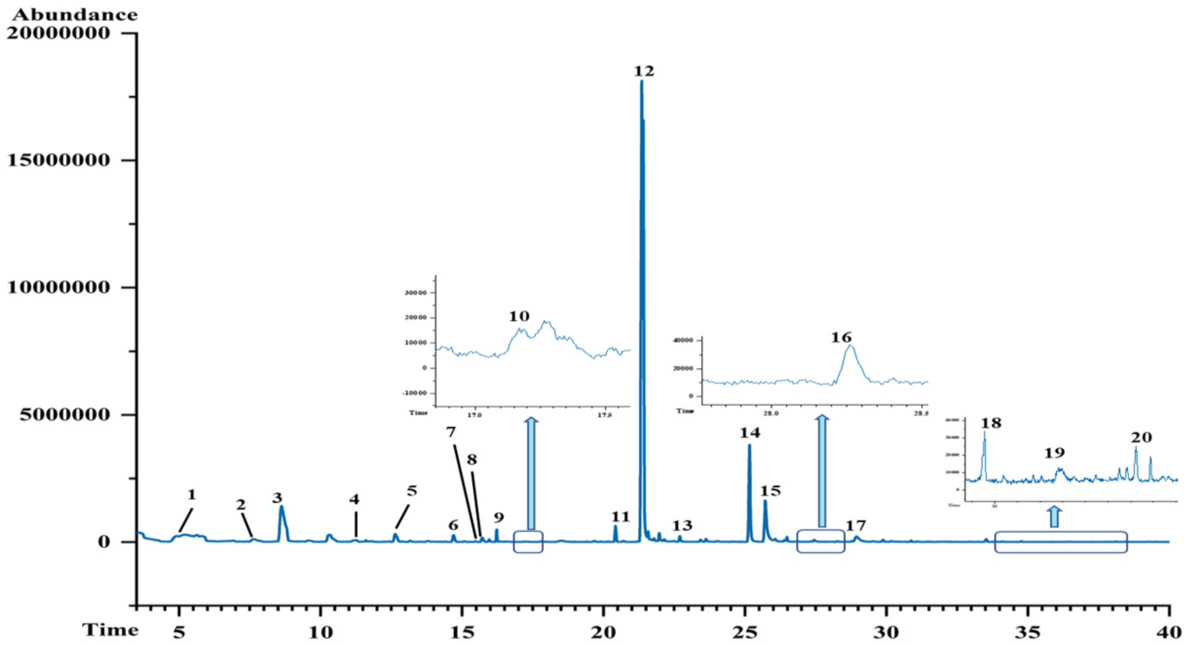


**Online resource 3: Gas chromatogram of a representative MF-HSVs identified from male fishing cat.**

**1. 2-Pentanone; 2. 2-Pentanone, 3-ethyl-; 3. 4-Heptanone; 4. 2-Butenal, 3-methyl-; 5. 3-Buten-1-ol, 3-methyl-; 6. 2-Hexanone, 3,4-dimethyl-; 7. 2-Buten-1-ol, 3-methyl-; 8. Thiophene, 2-methoxy-5-methyl-; 9. 1-Hexanol; 10. Nonanal; 11. 2-Ethyl-1-hexanol; 12. Benzaldehyde; 13. 3-Mercapto-3-methylbutyl formate (ester); 14. Acetophenone; 15. 3-Mercapto-3-methylbutanol; 16. 3-Methyl-3-methylthio-1-butanol; 17. Benzenemethanol, .alpha.-methyl-; 18. Phenol; 19. p-Cresol; 20. 3-Methyl-3-(2-methyldisulfanyl)-1-butanol.**

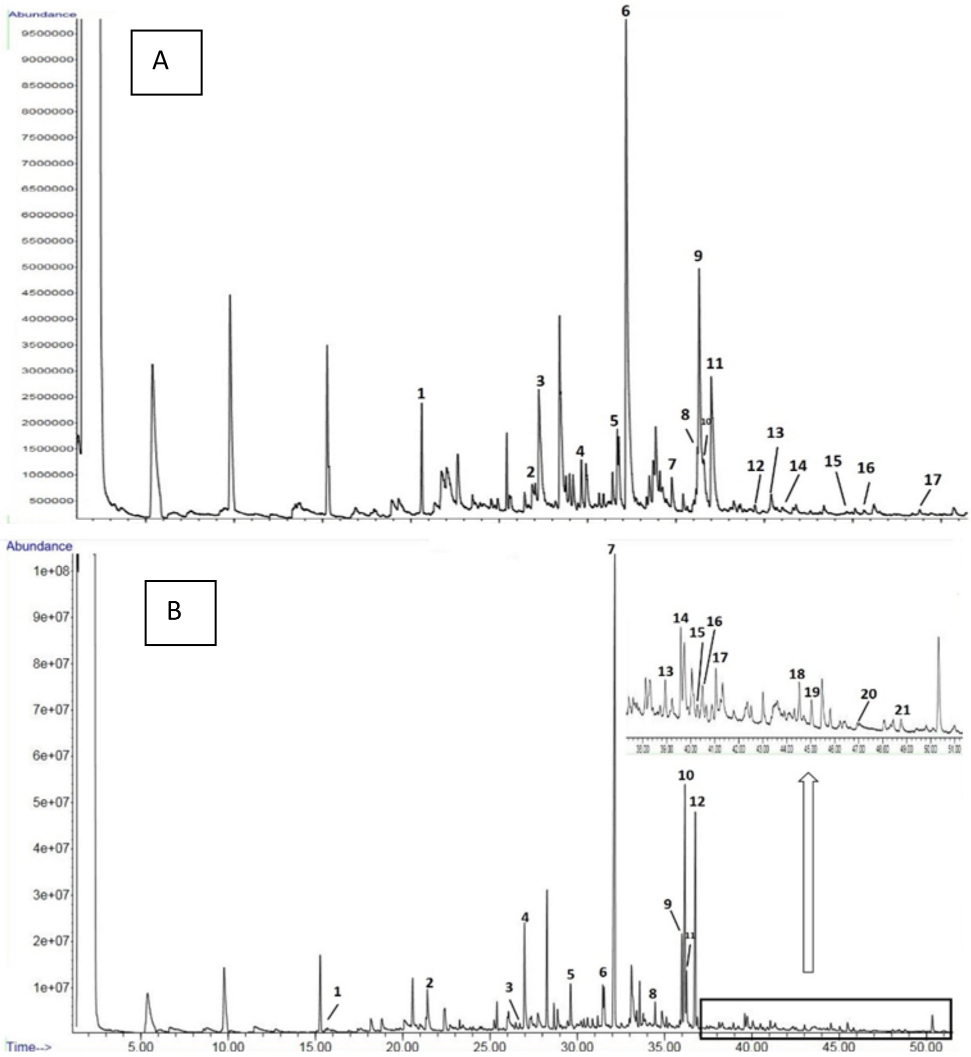


**Online resource 4: Gas chromatograms of FAs identified from UR (A) and MF (B) of Fishing cat.**

**A- 1. Dodecanoic acid methyl ester 2. cis-9-tetradecenoic acid methyl ester 3. Methyl tetradecanoate 4. Pentadecanoic acid methyl ester 5. 9-Hexadecenoic acid methyl ester 6. Hexadecenoic acid methyl ester 7. Heptadecanoic acid methyl ester 8. 9,12-Octadecadienoic acid methyl ester 9. 11-Octadecenoic acid methyl ester 10. 9-Octadecenoic acid methyl ester 11. Methyl stearate 12. Nonadecanoic acid methyl ester 13. 11-Eicosenoic acid methyl ester 14. Eicosanoic acid methyl ester 15. 13-Docosenoic acid methyl ester 16. Docosanoic acid methyl ester 17. Tetracosanoic acid methyl ester.**

**B- 1. Decanoic acid methyl ester 2. Dodecanoic acid methyl ester 3. cis-9-tetradecenoic acid methyl ester 4. Methyl tetradecanoate 5. Pentadecanoic acid methyl ester 6. 9-Hexadecenoic acid methyl ester 7. Hexadecenoic acid methyl ester 8. Heptadecanoic acid methyl ester 9. 9,12-Octadecadienoic acid methyl ester 10. 13-Octadecenoic acid methyl ester 11. 9-Octadecenoic acid methyl ester 12. Methyl stearate 13. Nonadecanoic acid methyl ester 14. 5,8,11-Eicosatrienoic acid, methyl ester 15. 11,14 Eicosadienoic acid methyl ester. 16. 11-Eicosenoic acid methyl ester 17. Eicosanoic acid methyl ester 18. 13-Docosenoic acid methyl ester 19. Docosanoic acid methyl ester 20. Tricosanoic acid methyl ester 21. Tetracosanoic acid methyl ester.**
